# Endothelial basement membrane laminins as an environmental cue in monocyte differentiation to macrophages

**DOI:** 10.1101/2020.04.15.043190

**Authors:** Lixia Li, Jian Song, Omar Chuquisana, Melanie-Jane Hannocks, Sophie Loismann, Thomas Vogl, Johannes Roth, Rupert Hallmann, Lydia Sorokin

## Abstract

Monocyte differentiation to macrophages is triggered by migration across the endothelial barrier, which is constituted by both endothelial cells and their underlying basement membrane. We address here the role of the endothelial basement membrane laminins (laminins 411 and 511) in this monocyte to macrophage switch. Chimeric mice carrying CX3CR1-GFP bone marrow were employed to track CCL2-induced monocyte extravasation in a cremaster muscle model using intravital microscopy, revealing faster extravasation in mice lacking endothelial laminin 511 (*Tek-cre::Lama5*^*-/-*^) and slower extravasation in mice lacking laminin 411 (*Lama4*^*-/-*^). CX3CR1-GFP^low^ extravasating monocytes were found to have a higher motility at laminin 511 low sites and to preferentially exist vessels at these sites. However, *in vitro* experiments reveal that this is not due to effects of laminin 511 on monocyte migration mode nor on the tightness of the endothelial barrier. Rather, using an intestinal macrophage replenishment model and *in vitro* differentiation studies we demonstrate that laminin 511 together with the attached endothelium collectively promote monocyte differentiation to macrophages. Macrophage differentiation is associated with a change in integrin profile, permitting differentiating macrophages to distinguish between laminin 511 high and low areas and to migrate preferentially across laminin 511 low sites. These studies highlight the endothelial basement membrane as a critical site for monocyte differentiation to macrophages, which may be relevant to the differentiation of other cells at vascular niches.

## Introduction

Monocytes are a diverse population that vary in origin, *in vivo* localization and function. They represent an important early line of defense against infectious agents or reaction to tissue damage and in several tissues bone marrow derived monocytes play a critical role in the replenishment of resident monocytes, macrophages and dendritic cells (DCs) in certain tissues, required for tissue immunity (Wynn et al., 2013). Trafficking and extravasation are, therefore, critical to their function. At least two subsets of monocytes exist, inflammatory monocytes that extravasate into tissues inflammatory conditions and the circulating monocytes, that can be distinguished by their expression of Ly6C, CCR2 and CX3CR1 (Gordon and Taylor, 2005, Auffray et al., 2007). However, data suggest that a continuum of monocyte phenotypes occur in vivo that are modulated by tissue/environmental factors (Dal-Secco et al., 2015). Several studies have demonstrated that the endothelium plays a role in this switch to macrophages (Jakubzick et al., 2013, Wynn et al., 2013); whether the underlying endothelial basement membrane also contributes to this step has not been considered to date.

Our previous work has shown that the basement membrane of vascular endothelium affects leukocyte extravasation by influencing adhesion and migration of extravasating cells (Sixt et al., 2001, Wu et al., 2009), and by affecting the tightness of endothelial monolayer (Song et al., 2017). Laminins were identified as the effector molecules in endothelial basement membranes, with laminin 511 (composed of α5, β1, γ1 chains) playing a decisive role. Endothelial basement membranes of arteries, arterioles and capillaries typically show high expression of the two main endothelial laminins, laminin 411 and 511, with progressively less laminin 511 occurring in the postcapillary venules, venules and veins, resulting in patches of laminin 411 positive and laminin 511 low expression which define the preferred sites of leukocyte extravasation (Sixt et al., 2001, Wang et al., 2006b, Wu et al., 2009). Although limited, data suggest that laminin 511 affects extravasation of different leukocyte types in different manners; while T cell migration is inhibited by direct interaction with laminin 511 (Wu et al., 2009), neutrophils can migrate across this substrate *in vitro* and their avoidance of laminin 511 high sites *in vivo* rather correlates with the higher endothelial-to-endothelial cell junctional adhesion strength at these sites (Song et al., 2017). Recent intravital imaging of sterile inflammation in the cremaster muscle also revealed that extravasating myeloid cells spend disproportionately long periods of time at the interface between the endothelial monolayer and the endothelial basement membranes and in breaching the basement membranes compared to their rapid penetration of the endothelial monolayer (Song et al., 2017), suggesting that the subendothelial compartment may be a site of reprogramming required for subsequent steps (Zhang et al., 2020).

Macrophages have been shown to bind and migrate on laminin *in vitro* (Mercurio and Shaw, 1988). While the laminin isoform employed in these studies does not occur in endothelial basement membranes, it shares similarities with endothelial laminin 511 and the receptors identified to be involved in macrophage-laminin interactions also recognize the endothelial laminin isoforms (Shaw et al., 1993). Macrophages, however, are not likely to extravasate across an endothelial basement membrane membrane *in vivo*, rather they are the consequence of monocyte extravasation or constitute a tissue resident population that forms during development. Due to the difficulty in obtaining sufficient numbers of monocytes for *in vitro* analyses, there is a paucity of information on monocyte-extracellular matrix interactions relevant to the extravasation process (Cougoule et al., 2012). We, therefore, here use live imaging of CCL-2 induced monocyte extravasation in the cremaster muscle model to investigate whether the endothelial basement membrane represents an environmental factor that affects monocyte phenotype and/or infiltration into tissues. Chimeric mice carrying CX3CR1-GFP bone marrow are used to track extravasating monocytic cells in WT host mice and in mice lacking the main endothelial basement membrane laminins, laminin 411 (*Lama4*^*-/-*^) and laminin 511 (*Tek-cre::Lama5*^*-/-*^). As reported for other leukocyte types, monocyte extravasation occurred preferentially at laminin 511 low sites; however, *in vitro* experiments revealed that this was not due to direct effects of laminin 511 on migration modes, nor effects on the tightness of the endothelial monolayer. Rather, using an intestinal macrophage replenishment model we demonstrate that laminin 511 together with the attached endothelium provide a decisive cue in promoting monocyte differentiation to macrophages, which in turn affects the extent of tissue infiltration.

## Materials and Methods

### Mice

Wild-type (WT) C57BL/6, laminin α4 knockout mice *(Lama4*^*-/-*^) (Thyboll et al., 2002), endothelial specific laminin α5 knockout mice (*Tek-cre::Lama5*^*-/-*^) (Song et al., 2013, Di Russo et al., 2017), and CX3CR1^GFP/+^ mice (Jung et al., 2000) were employed. Equal proportions of male and female mice were employed and no difference were noted between sexes. Animals were held in controlled temperature, humidity and day/night cycles according to the ARRIVE guidelines; female and pre-mating male mice were maintained in groups of a maximum of 5 animals/IIL cage, post-mating males were maintained in separate cages. All animal experiments were performed according to German Animal Welfare guidelines.

### Bone marrow chimera generation

Recipient *Lama4*^-/-^ and *Tek-cre::Lama5*^*-/-*^mice and their corresponding WT littermates were lethally irradiated at 15 Gy. 10^7^ bone marrow cells from femurs and tibias of CX3CR1^GFP/+^ mice were intravenously (i.v.) injected into each irradiated mouse. Reconstitution efficiency was analysed at 6 weeks for CD45^+^GFP^+^ versus CD45^+^GFP^-^ cells, or using the polymorphic lineage determinants (CD45.1 or CD45.2), for tracking GFP^+^ CD45.1^+^ donor cells in CD45.2 hosts. Only mice with >95% donor cell engraftments were employed for intravital microscopic imaging.

### Cells

Human umbilical venular endothelial cells (HUVECs) were cultured in endothelial cell growth medium (Promo Cell) at 37°C and 5% CO_2_ and used up to passage 5. Endothelioma cell lines were generated from *Lama4*^*-/-*^ embryos (eEND4.1) and WT littermates (eENDwt) as described previously (Williams et al., 1989); presence or absence of *Lama4* was checked by PCR and laminin α5 protein was quantified by Western blot. All endothelioma cell lines were cultured in DMEM plus 10% fetal calf serum (FCS) at 37°C and 5% CO_2_.

Murine Hoxb8 is a myeloid progenitor cell line that can be differentiated to monocytes or macrophages upon removal of β-estradiol (Sigma-Aldrich) (Wang et al., 2006a). Hoxb8 progenitor cell lines were generated from CD18^-/-^ C57BL/6 mice and their WT littermates (Scharffetter-Kochanek et al., 1998)and were cultured in RMPI 1640 containing 10% FCS, 40ng/ml GM-CSF (Promokine) and 1 μM β-estradiol. Differentiation to monocyte-like Hoxb8 cells was induced by removal of β-estradiol from the culture medium for 3 days (Wang et al., 2006a).

Bone marrow was flushed from tibias and femurs of WT C57/BL6 or CD18^-/-^ mice, sieved through a 70 μm sieve and red blood cells lysed in lysis buffer. The cells were pelleted and resuspended in DMEM with 10%FCS. The cells were incubated for 24 h at 37°C and 5% CO_2_ on bacterial plates, non-adherent cells were then harvested, transferred to a new bacterial plate in DMEM plus 10% FCS supplemented with 20 ng/ml M-CSF (Promokine) and cultured for 8 days to generate bone marrow derived macrophages (BMDMs).

Human monocytes were isolated from buffy coats of blood from healthy donors. A two-step procedure with a single gradient in each step was used, which included centrifugation through a Pancoll gradient (Pan-Biotech) (density = 1.070 g/ml) followed by a Percoll (General Electric Company) gradient (density = 1.064 g/ml). Monocyte purity was >85% as defined by flow cytometry for CD14 antibody (M5E2, Biolegend). Monocytes were cultured overnight in McCoy’s medium (Sigma-Aldrich) plus 15% FCS at 37°C and 7% CO_2_ in Teflon cell culture bags (OriGen) prior to use.

### Intravital microscopy

Mice were anesthetized using intra peritoneal (i.p.) injection of a mixture of ketamine hydrochloride (125mg/kg; Sanofi Winthrop Pharmaceuticals) and xylazine (12.5 mg/kg; TranquiVed, Phoenix Scientific) in saline. Intra-scrotal injection of 500 ng CCL2 (Biolegend) in 0.2 ml saline 3.5 h or 1.5 h before cremaster muscle exteriorization was employed to induce monocyte extravasation. Intravital microscopy was performed using an Axioscope A1 microscope equipped with an immersion objective (SW 40/0.75 NA) and stroboscopic epifluorescent illumination (Colibri 2, Zeiss), as described previously (Song et al., 2017). Blood flow center-line velocity was determined by imaging fluorescent beads (0.5μm, Polysciences) injected i.v. before image acquisition, which were converted to mean blood flow velocities by multiplying with an empirical factor of 0.625 (Lipowsky and Zweifach, 1978). Recorded images were analysed using ImageJ software. In some experiments, Alexa 650-labelled anti-laminin α5 (antibody 4G6) was injected intra-scrotally to mark the endothelial basement membrane (Song et al., 2017); intravital microscopy was otherwise performed as described above.

To correlate speeds of migration with CX3CR1-GFP expression levels, the ImageJ plugin-TrackMate was used to track the motility of CX3CR1-GFP^high^ and CX3CR1- GFP^low^ cells in laminin 511^high^ and laminin 511^low^ areas; LoG detector was used to detect the CX3CR1-GFP cells and Simple LAP tracker was used for tracking the cells. For speed analyses, at least 60 frames were selected for each track and the mean speed of the track was calculated. Each tracked cell was categorized as CX3CR1- GFP^high^ or CX3CR1-GFP^low^, based on the initial CX3CR1-GFP^GFP^ fluorescent intensity.

Experiments were performed at least 3 times/chimera. In all experiments 3-9 different postcapillary venules were imaged/ mouse and analysed. Experiments were performed using age and sex matched WT chimera and laminin knockout chimeras. Biological replicates were individual animals/ chimeras, technical replicates were analyses performed on different postcapillary venules in the same animal.

### Confocal microscopy

After live cell imaging the cremaster muscle was removed, fixed in 1% PFA for 1h, blocked and permeabilized in PBS containing 5% FCS and 0.5% Triton X-100 for 30 min. Incubations with primary antibodies were performed overnight at 4°C, followed by secondary antibody incubation for 2 h at room temperature. To quantify the proportions of CX3CR1-GFP+MHCII+ cells in total GFP+ cells, the cremaster muscles were stained by using primary antibodies including rabbit anti-mouse laminin α5 (antibody 405) or anti-mouse laminin α4 (antibody 377) and Brilliant Violet 411-conjugated rat anti-mouse MHC II (Biolegend); monocytes were identified by CX3CR1^GFP^ expression. Alexa 647- (Dianova) anti-rabbit secondary antibodies were employed. The 2-7 postcapillary venules were imaged/ chimera and the images were analysed using ImageJ software.

Dynamic *in situ* cytometry (Bousso and Moreau, 2012) was employed to investigate associations between laminin 511 expression and CX3CR1-GFP mean fluorescence intensity (MFI): cremaster muscles were stained with rat anti-mouse PECAM-1 (MEC13.3, BD Bioscience) and rabbit anti-mouse laminin α5 (antibody 405) (Sixt et al., 2001) and monocytes were identified by CX3CR1^GFP^ expression; Alexa 647- conjugated anti-rat and Alexa 555-conjugated anti-rabbit secondary antibodies (Abcam) were employed. CX3CR1-GFP expression was analysed with Flowjo v10.4 and the image files were converted into ‘flow cytometry standard’ files for analysis by flow cytometry software. Image pixels were reconstituted relative to locations in the X and Y axes; laminin 511^high^ and laminin 511^low^ regions were gated separately and CX3CR1-GFP MFI were analysed in these two regions. Data were expressed as percent CX3CR1-GFP^low^ cells of total CX3CR1-GFP positive cells in laminin 511^high^ and laminin 511^low^ regions. These analyses were performed on 5 postcapillary venules in each of 3 mice/chimera.

### Adhesion assay

Cell attachment assays were performed using dot adhesion assays. 12-well Nunc Maxisorb plates were coated overnight at 4°C with 5 μl dots of 20 μg/ml laminin 411, laminin 511, VCAM-1 (R&D) or ICAM-1 (R&D), wells were blocked with 1% BSA for 1 h at 37 °C and washed with PBS before use. Laminins 411 and 511 were purified as described previously (15). 1 × 10^6^ cells in adhesion buffer (DMEM, 0.5% BSA, 50 mM HEPES, pH 7.5) were added to coated wells and allowed to adhere for 1 h at 37°C. Non-adherent cells were removed and attached cells were fixed in 4% paraformaldehyde (PFA) for 10 min and stained by 0.5% crystal violet for 10 min. Cells bound to the dots of different substrates were imaged using an inverse Axiovert microscope (Zeiss) and the number of adherent cells per substrate was quantified using ImageJ software. Adhesion assays were carried out using human monocytes and mouse monocyte-like Hoxb8 cells.

Inhibition assays to assess receptors responsible for adhesion to different substrates involved pre-incubation of cells at 4°C for 30 min prior to addition to protein-coated 12- well plates with 20 μg/ml anti-human integrin β2 (antibody TS1/18), anti-human integrin β1 (antibody P5D2) and anti-mouse integrin β3 (antibody 2C9.G2), or 50 μg/ml anti- mouse integrin β1 (antibody Ha2/5) and anti-human and mouse integrin α6 (antibody GoH3). Inhibition assays were performed using 20 μg/ml of extracellular matrix protein.

Biological replicates were different batches of Hoxb8 cells, human monocytes or BMDM preparations used in separate experiments, technical replicates were repetitions of the same test (eg adhesion on a specific laminin isoform) in one experiment.

### Migration assays

24-well Transwell inserts (6.5 mm^2^) with polycarbonate filters (5 μm pore size) (Corning Costar) were used for transmigration assays. Membranes were coated overnight at 4 °C with 10 μg/ml laminins 411 or 511 and blocked with 1% BSA in PBS for 1 h at 37°C. Human monocytes, mouse Hoxb8 monocytes and BMDM were used in transmigration assays. 5 × 10^5^ cells in 100 μl adhesion buffer were added to the upper chamber and either 500 ng/ml CCL2 (R&D Systems) or 10 nM C5a (R&D Systems) was added to the bottom chamber. Cells were permitted to transmigrate at 37°C for 4h, and the number of transmigrated cells was expressed as a percentage of the total cells initially added. For trans-endothelial migration, HUVEC or bEND.5 cells were plated at confluent densities onto the laminin 411, 511 or 111 coated filters and stimulated for 4 h with 5nM tumor necrosis factor-α (TNF-α; R&D Systems), then human monocytes or monocyte Hoxb8 cells were added. These experiments were 3 times with triplicates/ experiment.

2D Chemotaxis assays utilized the Ibidi µ-Slide Chemotaxis system and 20nm C5a as chemoattractant. Slides were coated overnight at 4°C with 10 μg/ml laminin 411 or 511, and mouse WT or CD18^-/-^ lipopolysaccharide (LPS/1 µg/ml) (Sigma) activated BMDM cells (1×10^6^ cells/ml) were added in attachment buffer (RPMI 1640 medium, 20 mM Hepes, 10 % FCS) and monitored by time-lapse video microscopy every 2 min for 24h. Cell migration tracks between 6-12 h were analysed by ImageJ using a manual tracking plugin and the Ibidi software chemotaxis and migration tool. A minimum of 20 randomly selected cells on each substrate were manually tracked and cell velocities were measured/ experiment. The experiment was repeated 3 times.

### Isolation of colonic lamina propria (LP) cells

LP cells were purified from colons of adult (8-12 week-old) and day 21 postnatal (P21) mice by enzymatic digestion (Bain et al., 2014). Colons were excised and washed in PBS, opened longitudinally and cut into 0.5 cm sections, and shaken vigorously in Hank’s balanced salt solution (HBSS) containing 2% FCS. To remove the epithelial layer, 2 mM EDTA in Ca^2+^/Mg^2+^ free HBSS was added twice with shaking for 15 min each at 37°C. The remaining tissue was incubated in digestion buffer (RPMI 1640, 10% FCS, 1.25 mg/ml collagenase D (Roche), 0.85 mg/ml collagenase V (Sigma– Aldrich), 1 mg Dispase (Gibco), and 30 U/ml DNase (Roche)) for 30–45 min at 37°C with shaking. The resulting cell suspension was passed through a 40 μm cell strainer and washed twice in RPMI 1640. The experiments for adult and P21 mice were done 3 times with at least 2 mice/mouse strain/experiment. Typically, 4-5 million single cells were obtained from 1 adult colon, and approximately 2 million single cells from one P21. Total numbers of cells isolated in adult or P21 colons were the same regardless of genotype.

### Flow cytometry

Colonic LP samples (2 million cell/staining) were stained at 4°C for 20 min with the following antibodies: Ly6C-FITC (AL-21, BD Pharmingen), F4/80-PE (BM8, BioLegend), Siglec F-PE-CF594 (E50-2440, BD Bioscience), Ly6G PE-Cy7 (1A8, BioLegend), CD11b-APC (M1/70, eBioscience), CD11c-AF700 or -APC-R700 (N418, BioLegend), MHCII-BV421 (M5/114.15.2, BioLegend), and CD45-BV510 (30-F11, BioLegend)) at 4°C for 20 min. Dead cells were identified by staining with eFluor 780 Fixable Viability dye (eBioscience) and gatings were performed as described previously (Bain et al., 2014).

For analyses of integrin receptors on human monocytes, mouse Hoxb8 monocytes and mouse BMDMs, antibodies to integrins β1 (mouse 9EG7 (Lenter et al., 1993)/ human P5D5) (Wayner et al., 1988), α6 (GoH3, BD Pharmingen), α5 (mouse 5H10-27, BD Pharmingen / human P1D6) (Wayner et al., 1988), α4 (PS/2, Abcam), α3 (mouse polyclonal, R&D/ human P1B5) (Wayner et al., 1988), β3 (mouse 2C9.G2, Biolegend/ human B3A, Chemicon) and huan β2 (mouse C71/16, BD Pharmingen / human TS1/18, Thermo Scientific) were employed. Cells were analysed with a Gallios (Beckman Coulter) or Celesta (BD) flow cytometer and FlowJo software.

### *In vitro* monocyte differentiation

3D Collagen type I was prepared using 8 parts rat collagen type I (Sigma) and 1 part 10x DMEM and was neutralized to pH 7.0-7.5 by adding 0.5 parts 0.5N NaOH; 400 µl collagen I gel was then added immediately to each well of a 24-well plate. Endothelial cells (e.END4.1, e.ENDwt) were seeded onto the collagen type I gel base and were incubated at 37°C overnight. Splenic monocytes were isolated by depleting CD19, Ly6G, CD11c, CD3ε, Siglec-F, MHCII and CD11C positive cells using magnetic beads (Miltenyi Biotec); 5 ×10^5^ cells were plated onto the confluent endothelial cells for 16 h at 37°C. The conditioned medium was collected for cytokines analyses, and remaining unattached cells were removed by 5x washing with DMEM; 400 µl Collagenase D (2 mg/ml, Sigma) was then added and the cultures digested at 37°C for 30 min. Cells were harvested and analysed by flow cytometry for PECAM-1, Ly6C, MHCII, CD11b and F4/80 expression. The same experiments were performed with splenocytes preincubated for 30 min in the presence of 25 μg /ml integrin α6 function blocking antibody (GoH3) or an isotype control. Experiments were performed 3 times using cells from 2 mice. Cytokines were measured in 5x concentrated and conditioned media using Th1/Th2 10 Plex Flowcytomix Kit (eBioscience).

### Statistical analyses

Data sets were tested for normal distribution with a D’Agostino & Pearson normality test. If all data sets of one experiment passed the normality test, significance was analysed with a Welch’s t-test. A Mann-Whitney test was applied if the data was not normally distributed. P-values of <0.05 were considered significant.

## Results

### Monocyte Extravasation in the Cremaster Muscle Model

Bone marrow chimeras were generated by transferring bone marrow cells from CX3CR1^GFP/+^ transgenic mice to *Lama4*^-/-^ and *Tek-cre::Lama5*^-/-^ hosts and their corresponding wild type (WT) littermates; the extravasation specifically of monocytes was induced by intrascrotal injection of CCL2. Postcapillary venules in the cremaster muscle were intravitally imaged for 2 h or 4 h and the extent of monocyte extravasation was determined by quantification of GFP^+^ cells in an area extending 75 μm from each side of a vessel over a distance of 100 μm vessel length (representing 1.5 × 10^4^ μm^2^ tissue area); GFP+ cells adherent to the vessel lumen for 30 sec in a 307μm length were counted and expressed relative to the surface area of the cylinder vessel tube (Fig. 1A, Fig. S1, Movie 1) (Voisin et al., 2009, Song et al., 2017).

**Fig. 1.**
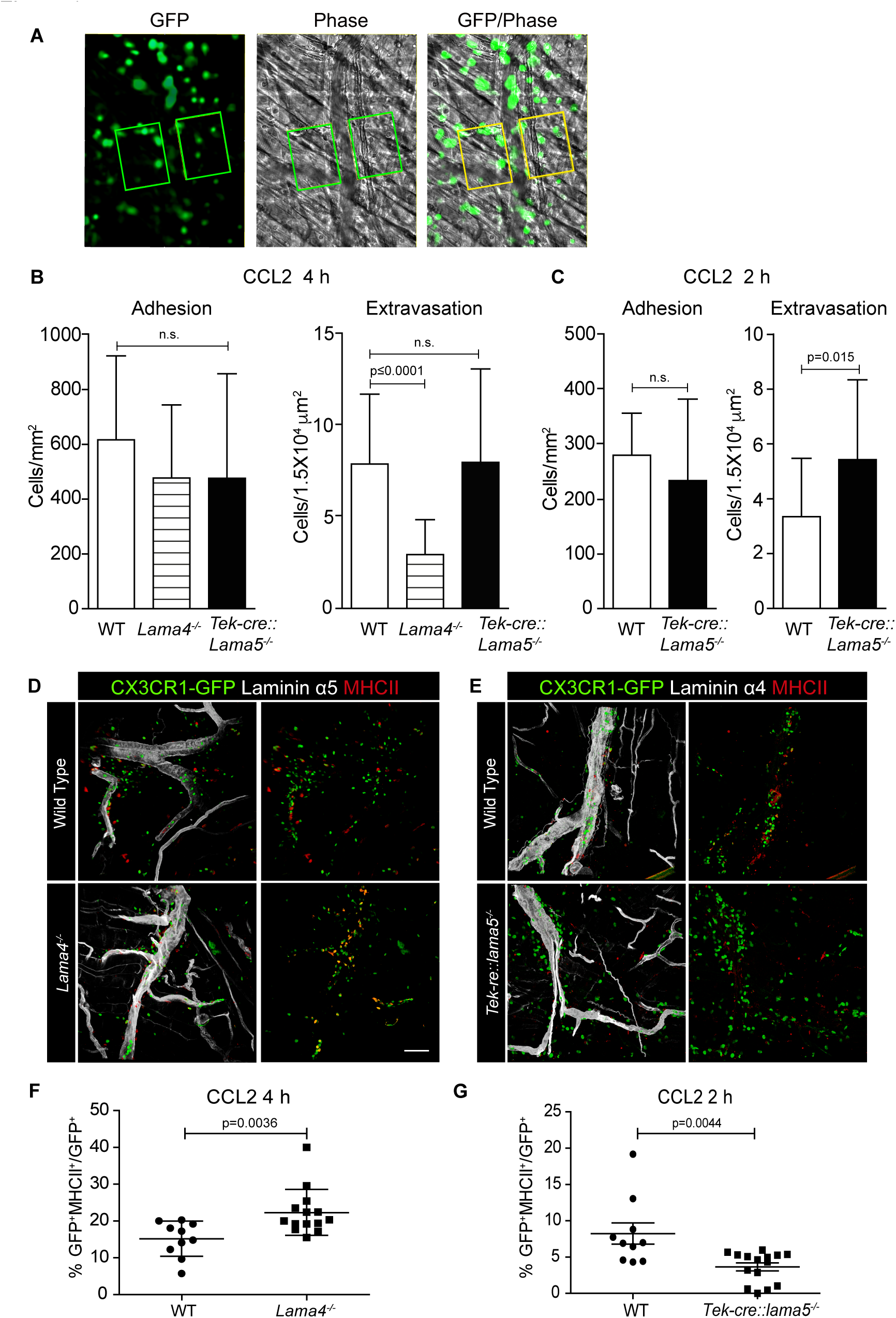
Intravital microscopy of CCL-2 induced monocyte extravasation in cremaster muscle postcapillary venules of *Lama4*^*-/-*^ or Tek-cre::*Lama5*^*-/-*^ hosts carrying CX3CR1^GFP/+^ bone marrow. (A) Representative fluorescence and corresponding phase contrast images from movies of CX3CR1-GFP^+^ cells extravasating across postcapillary venules of the cremaster muscle; boxed areas are 75 μm from either side of the postcapillary venule for a length of 100 μm, used to quantify extravasated CX3CR1-GFP^+^ cells; scale bars = 50μm. Quantification of adherent and extravasating CX3CR1-GFP^+^ monocytes at (B) 4 h and/or (C) 2 h post-application of CCL2. Data are means ± SD from 5 WT mice and 4 mice for each chimera with 4-8 postcapillary venules examined/mouse at 4h and 3-4 WT and chimeric mice with 3-4 postcapillary venules examined/mouse at 2h. (D,E) Excised tissues were immunofluorescently stained for MCHII and laminin α5 or α4 to identify CX3CR1-GFP^+^MCHII^+^ extravasating cells in relation to the endothelial basement membrane; scale bar = 50μm. (F, G) Quantification of proportions of CX3CR1- GFP^+^MCHII^+^ cells expressed as percentage of total GFP^+^ cells in *Lama4* ^*-/-*^ or Tek- cre::*Lama5*^*-/-*^ hosts. Data are means ± SD from 4 WT and 4 chimeric mice with 2-4 postcapillary venules examined/mouse in (F) and 3 WT and 3 chimeric mice with 3-7 postcapillary venules examined/mouse in (G). Significance in B, C, F and G were calculated using Mann-Whitney tests as not all data passed the normality test by D’Agostino & Pearson; the exact *p*-values are shown in the graphs.

The number of adherent GFP^+^ monocytes in postcapillary venules were similar in the three chimeras (Fig.1 B, C). However, the number of extravasated GFP^+^ monocytes at 4h after CCL2 application was significantly lower in *Lama4*^-/-^ hosts compared to WT hosts (Fig. 1 B). By contrast, significantly higher numbers of GFP+ monocytes had extravasated at 2h in *Tek-cre::Lama5*^-/-^ hosts compared to WT littermates, but this difference was no longer detectable at 4h (Fig. 1B,C), suggesting that monocytes migrated faster across postcapillary venules that lack laminin α5. To more precisely identify extravasated cells, tissues were excised following intravital imaging and examined by confocal microscopy for expression of both GFP and MHCII which is only induced upon extravasation (Jakubzick et al., 2013); staining for endothelial basement membrane permitted localization of the monocytes within or outside vessels. This revealed a higher proportion of GFP^+^MHCII^+^ double positive cells in *Lama4*^-/-^ hosts which, however, remained associated with laminin 511 positive vessels compared to WT controls (Fig. 1D,F) and *Tek-cre::Lama5*^-/-^ hosts (Fig. 1E,G). In *Tek-cre::Lama5*^-/-^ hosts most GFP^+^ were located outside of the vessels and showed minimal MCHII expression (Fig. 1E,G), suggesting that even though more cells extravasated their differentiation towards macrophages was hampered.

### Monocyte Accumulation at Laminin α5^high^ Sites

As reported for other leukocytes (Wang et al., 2006b, Wu et al., 2009, Song et al., 2017), extravasating CX3CR1-GFP^+^ cells localized preferentially at laminin α5^low^ sites in WT hosts (Fig. 2A); however, extensive accumulation of CX3CR1-GFP^+^ cells also occurred within laminin α5^high^ vessels (Fig. 2A), which was not observed in TNF-α induced neutrophil extravasation (Song et al., 2017) probably because of their faster extravasation compared to monocytes and, hence, absence of accumulation within vessel lumens.

**Fig. 2.**
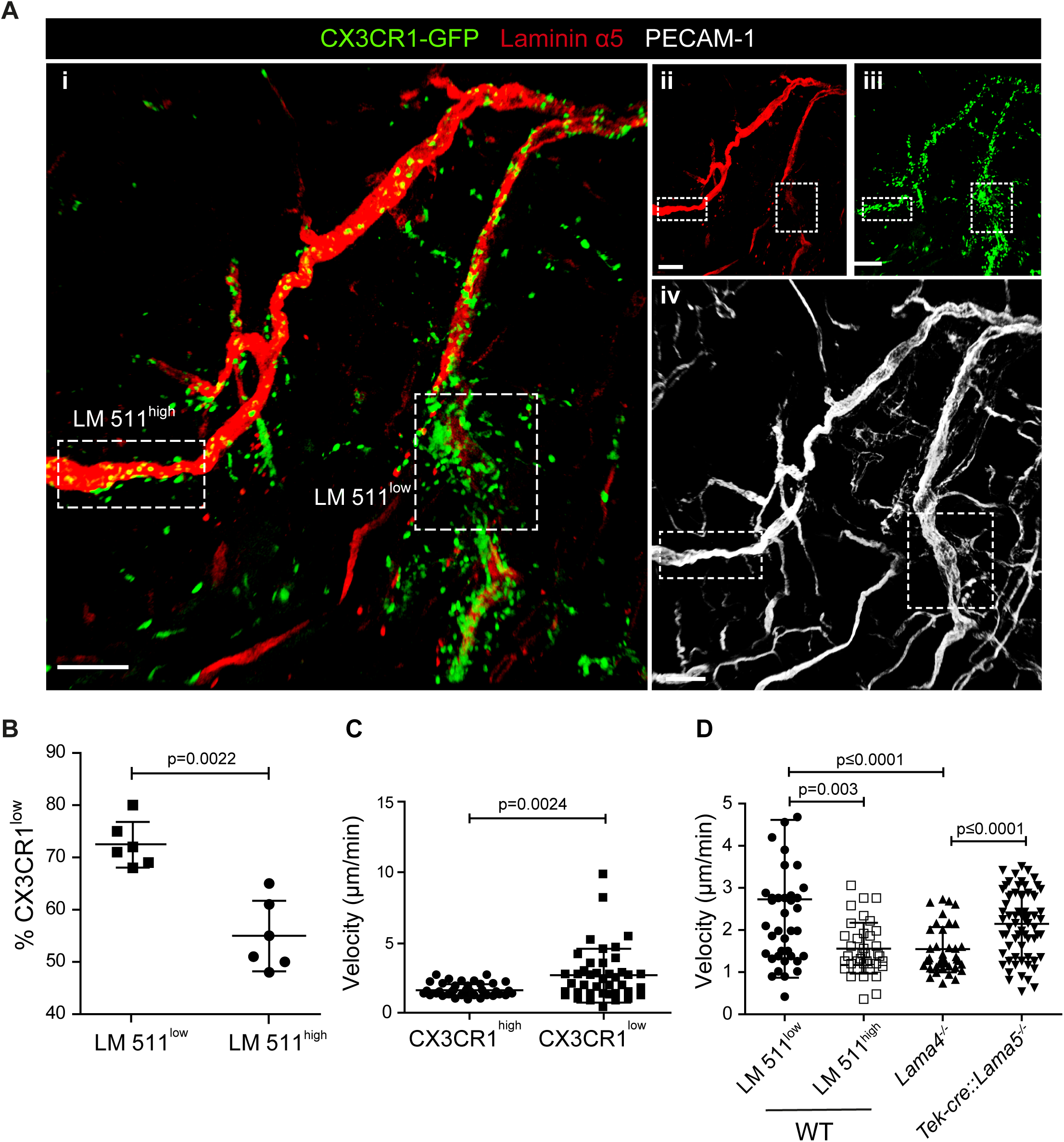
CX3CR1-GFP^low^ inflammatory monocytes preferentially extravasate at laminin 511^low^ sites. (A) Representative confocal microscopy of whole mount cremaster muscle from a WT host carrying CX3CR1^GFP/+^ bone marrow stained with anti-laminin α5 and anti-PECAM-1 (i); single channel images are shown in (ii, iii, iv); scale bar = 100 μm. Experiment was repeated 3 times with 1 animal with the same results. (B) To track CX3CR1-GFP^low^ inflammatory monocytes GFP mean fluorescence intensity (MFI) was measured *in situ* in areas of high and low laminin 511 expression (see Fig. S2A), revealing a higher proportion of CX3CR1-GFP^low^ cells, expressed as a percent of total GFP^+^ cells, at laminin 511^low^ compared to laminin 511^high^ sites; data are means ± SD of 3 mice from 3 separate experiments and at least 5 sites examined/mouse. (C) Quantification of GFP MFI and migration speed of individual CX3CR1^hi^ and CX3CR1^low^ cells analysed in 3 WT hosts from 3 separate experiments. (D) Average migration velocities of extravasating GFP^+^ cells at laminin 511 low and high sites in WT controls, and in *Lama4*^*-/-*^ and Tek-cre::*Lama5*^*-/-*^ chimeras. Data in C and D are means ± SD calculated from 39-64 cells analysed in 3 mice per chimera; movies were captured at 4h post-administration of CCL2. Significance of data in B, C and D were calculated using Mann-Whitney tests since the data did not pass the normality test by D’Agostino & Pearson; the exact *p*-values are shown in the graph.

As CX3CR1-GFP^low^ monocytes have been reported to mark extravasating inflammatory monocytes (Gordon and Taylor, 2005, Auffray et al., 2007), we investigated CX3CR1-GFP mean fluorescence intensity (MFI) in relation to laminin α5 high and low regions. CCL-2 induced extravasation was performed in chimeric mice in which Alex 550-labelled anti-laminin α5 was injected intra-scrotally to visualize laminin α5 (Song et al., 2017) (Movie 2) (Gordon and Taylor, 2005). CX3CR1-GFP MFI was measured in areas of high and low laminin 511 expression using dynamic *in situ* cytometry, confirming a higher proportion of CX3CR1-GFP^low^ cells at laminin 511^low^ than at laminin 511^high^ sites, and correspondingly higher proportions of CX3CR1- GFP^high^ monocytes at laminin 511^high^ sites (Fig. 2B, Fig. S2A). Correlation of CX3CR1- GFP MFI with speeds of cell migration, revealed that CX3CR1-GFP^low^ inflammatory monocytes had higher average speeds of migration than CX3CR1-GFP^high^ monocytes (Fig. S2B; Fig. 2C), as expected for extravasating cells. In *Lama4*^*-/-*^ hosts, which show a compensatory ubiquitous expression of laminin 511 in all endothelial basement membranes and the absence of laminin 511 low patches at postcapillary venules (Wu et al., 2009), all CX3CR1-GFP+ cells showed migration speeds similar to those measured for monocytes at laminin 511^high^ sites in WT hosts, while in *Tek-cre::Lama5*^*-*^ */-* hosts CX3CR1-GFP+ cells showed higher average migration velocities (Fig. 2D). Taken together, this suggests that the more motile CX3CR1-GFP^low^ inflammatory monocytes preferentially extravasate at laminin 511^low^ sites.

### Mouse monocyte adhesion on laminins *in vitro*

Monocytic cell lines (U-937, THP-1, Mono Mac 6) have been previously reported to bind to endothelial laminins (Pedraza et al., 2000) and macrophages have been reported to bind and migrate on the non-endothelial laminin 111 (Mercurio and Shaw, 1988). However, a comprehensive comparison of monocyte and macrophage binding to the endothelial laminins, laminin 411 and 511, and receptor involvement has not been carried out. As it is not possible to obtain sufficient mouse primary CX3CR1-GFP monocytes for *in vitro* adhesion and migration assays, mouse monocyte-like Hoxb8 cells (Wang et al., 2006a) and primary human monocytes were compared to bone marrow derived mouse macrophages (BMDM). Control substrates included ICAM-1 and VCAM-1, which are recognized by αLβ2 (LFA-1) and α4β1, respectively. Human monocytes (Fig. 3A), mouse Hoxb8 monocytes (Fig. 3B) and mouse BMDM (Fig. 3C) showed extensive binding to laminin 511, ICAM-1 and VCAM-1, and lower binding to laminin 411 (Fig. 3A, B). However, human monocytes and mouse Hoxb8 monocytes also showed high levels of binding to BSA, probably due to activation of integrin β2, which we have previously shown indiscriminately enhances non-specific binding of neutrophils to all substrates including BSA and plastic (Sixt et al., 2001b) (Fig. 3A, B). Therefore, mouse integrin β2 null (CD18^-/-^) (Scharffetter-Kochanek et al., 1998) Hoxb8 monocytes were generated and employed in all subsequent adhesion assays, and human monocytes were treated with anti-human integrin β2 blocking antibody (TS1/18). This significantly reduced binding to ICAM-1 and BSA, and revealed specific binding to laminin 511 and VCAM-1 for all cells (Fig. 3A-C); only mouse BMDM also showed binding to laminin 411 (Fig. 3C).

**Fig. 3.**
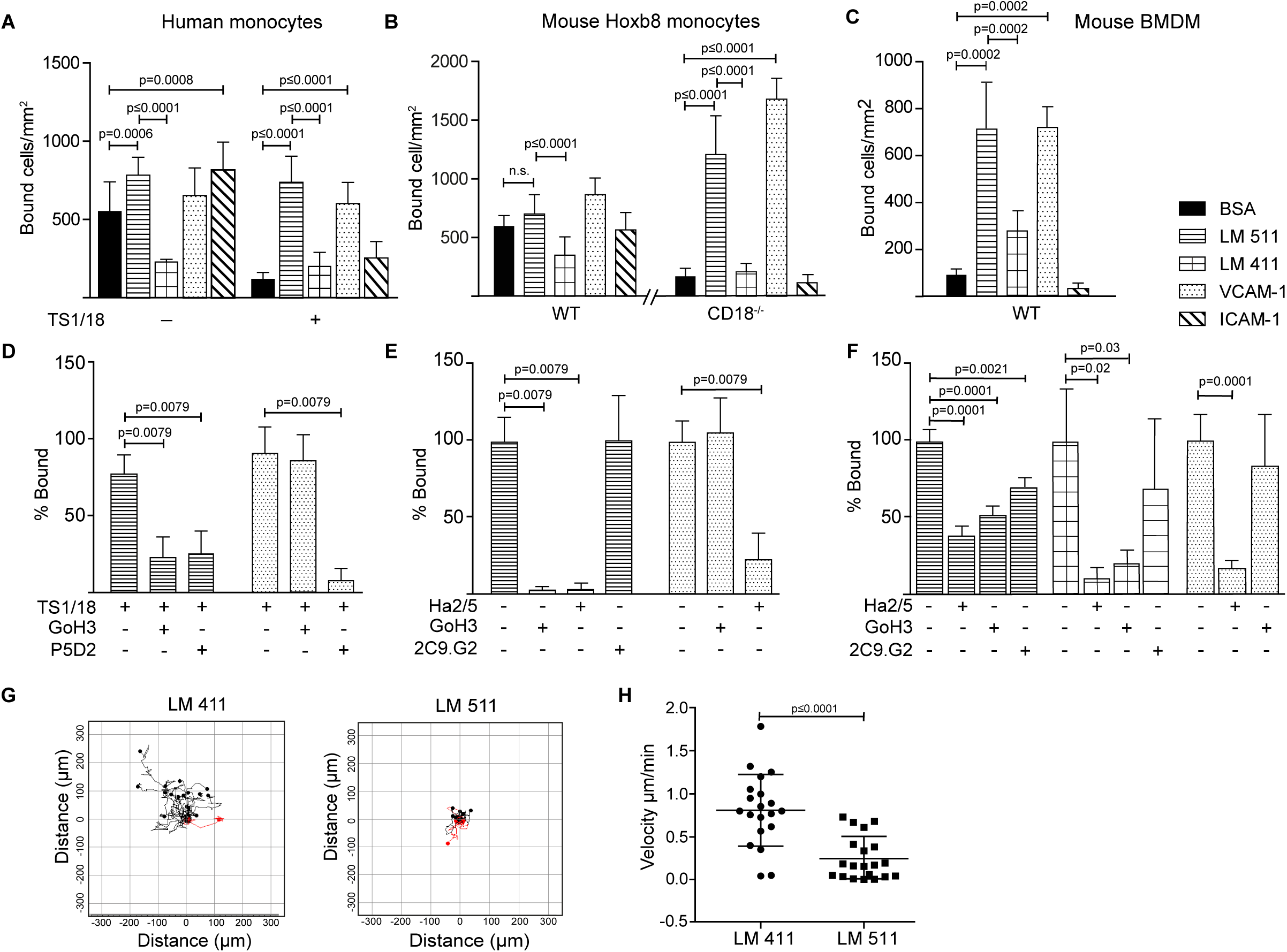
Human monocytes, mouse monocyte-like Hoxb8 cells and mouse bone marrow derived macrophages (BMDM) bind differentially to endothelial laminins. Human monocytes (A, D), mouse monocyte-like Hoxb8 cells (B, E) and BMDM (C, F) adhesion to laminins 411, 511, VCAM-1, ICAM-1 or BSA. To exclude non-specific integrin β2-mediated adhesion of monocytic cells, experiments were performed with untreated or integrin β2 antibody (TS1/18) treated human monocytes, and WT and CD18^-/-^ mouse monocyte-like Hoxb8 cell. Adherent cells were counted at 1 h and expressed as number of bound cells/mm^2^. (D) Inhibition assays were performed with human monocytes preincubated with TS1/18 and either anti-integrin α6 (GoH3) or β1 (P5D2) antibodies before addition to substrates. Similar experiments were performed with mouse monocyte-like CD18^-/-^ Hoxb8 cells (E) and MBDM (F) pre-incubated with anti-mouse integrin β1 (Ha2/5), α6 (GoH3) or β3 (2C9.G2) antibodies. Data in D-F are expressed as percent of total cells bound in the absence of blocking antibodies. All data except F are means ± SD from 4-5 independent experiments with triplicates/experiment; F are means ± SEM of 3 experiments with triplicates/experiment. (G) Random migration of BMDMs on laminin 411 and laminin 511 were imaged by time-lapse video microscopy; >20 cells on each substrate were tracked and analysed. (H) Velocity of migrating cells on each substrate; data are means ± SD from a representative experiment. The experiment was performed 3 times with similar results. Significance was calculated using a Mann-Whitney test in A,C,D,E and H, a Welch’s t-test was applied in B, and an unpaired t-test in F; exact *p*-values are shown in the graph.

Flow cytometry revealed similar expression patterns for the main laminin binding integrins on WT and CD18^-/-^ mouse Hoxb8 and human monocytes and BMDM which suggested expression of integrins α6β1, α5β1 and αvβ1 and/or αvβ3 (Fig. S3), with higher levels of α6β1, αvβ1 and αvβ3 and the unique expression of integrin α3β1 on BMDM (Kikkawa et al., 2000, Sasaki and Timpl, 2001, Nishiuchi et al., 2006). To test integrin involvement in binding to the laminins, function blocking antibodies to integrins α6, β1 or β3 were employed in adhesion assays, revealing complete inhibition of mouse Hoxb8 and human monocytes binding to laminin 511 by anti-integrin β1 and α6 antibodies (Fig. 3D, E). In the case of BMDM, anti-integrin β1 and α6 antibodies also ablated binding to laminin 411 but only partially inhibited binding to laminin 511, suggesting the involvement of other receptors (Fig. 3F). Partial inhibition by anti- integrin β3 suggests the involvement of αvβ3 (Fig. 3F), although αvβ1 cannot be excluded. While integrin α3β1 can also bind to laminin 511 (Kikkawa et al., 2000, Nishiuchi et al., 2006), the absence of function blocking antibodies against the mouse protein prevented investigation of whether this is the case in BMDM. This suggests that α6β1 is the major laminin 511 receptor on mouse and human monocytes (Sasaki and Timpl, 2001) while BMDMs also employ an αv series integrin, and potentially also α3β1, to bind laminin 511 and, additionally, bind to laminin 411 via integrin α6β1.

### Migration Assays

We have previously shown that splenic monocytes can equally well migrated across laminin 411, 511 or non-endothelial laminin 111 in a transwell assay (Wu et al., 2009). This pattern was not altered by plating endothelial cells onto the laminin substrates (Fig. S4A, B), as we have shown occurs with other immune cell types (Song et al., 2017), nor did the mouse HoxB8 monocytes, human monocytes or mouse BMDM employed here express laminin proteins (Fig. S5) which has been proposed by others to affect the extravasation process (Pedraza et al., 2000). However, BMDM migration on laminin 411 and 511 did differ: As macrophages are unlikely to transmigrate across endothelial monolayers and their basement membranes *in vivo*, we analysed 2D chemotaxis of BMDM using time-lapse video microscopy, revealing BMDM migration on both laminins 411 and 511 but more extensive and faster migration on laminin 411 compared to laminin 511 (Fig. 3G, H). Hence, even though macrophages in general migrate more slowly than other leukocytes (Bzymek et al., 2016), they can distinguish between laminins 411 and 511 and migrate differently on these substrates.

### Monocyte *in vivo* differentiation

As monocytes and macrophages have different migration capacities on different substrates (Mercurio and Shaw, 1988, Shaw et al., 1993, Shaw and Mercurio, 1994) and monocytes have been reported to initiate gene expression changes towards a more differentiated MCHII^+^ phenotype upon extravasation (Thomas-Ecker et al., 2007, Jakubzick et al., 2013), we investigated the possibility of *in vivo* modulation of the monocyte phenotype during extravasation process.

As it is not possible to isolate sufficient leukocytes from the cremaster muscle to phenotype the extravasated GFP^+^ cells, we investigated the intestine where resident macrophages of the lamina propria (LP) are constantly replenished by circulating monocytes (Zigmond and Jung, 2013). Differentiating monocytes and macrophages, therefore, represent a large fraction of the total LP cells and are well characterized by their differential expression of cell surface markers (Fig. S6A). This permits categorization into early extravasated Ly6C^high^MHCII^low^ immature P1 and maturing Ly6C^high^MHCII^high^ P2 populations, and mature Ly6C^low^MHCII^high^ P3 macrophages (Bain et al., 2014).

Postcapillary venules of the colon wall showed the same the patchy laminin α5 and even laminin α4 distribution as occurs in other tissues (not shown). LP cells from colons of *Lama4*^*-/-*^ and *Tek-cre::Lama5*^*-/-*^ mice and WT controls were analysed by flow cytometry, using the gating strategy outlined in Fig. S6A, at adult stages (3 month-old) (Fig. 4A-C) and at post-natal day 21 (P21) (Fig. 4D-E; Fig. S6B) when the peak of colonization of intestinal macrophages by circulating monocytes occurs. In P21 samples, P1 and P2 early extravasating populations were analysed together due to the lower numbers of cells obtained (Fig. S6B). In both adult and P21 LP, fewer early extravasating P1 and P2 cells were measured in *Lama4*^*-/-*^ and more in *Tek-cre::Lama5*^*-*^ */-* compared to WT mice (Fig. 4B, D), reflecting the same pattern of results that were observed in the intravital imaging studies in the cremaster muscle (Fig. 1A, B). However, despite fewer extravasating monocytes, a larger P3 mature macrophage population was measured in *Lama4*^*-/-*^ LP than in either WT or *Tek-cre::Lama5*^*-/-*^ LP as reflected by higher P3/P2 ratios (Fig. 4C, E). By contrast, *Tek-cre::Lama5*^*-/-*^ LP showed lower proportions of P3/P2 ratios than *Lama4*^*-/-*^ and WT LP (Fig. 4C, E), which was particularly evident in P21 LP (Fig. 4E, Fig. S6B) where a larger proportion of macrophages are derived from infiltrating monocytes (Bain et al., 2014). These results suggest that in the absence of endothelial basement membrane laminin 511, monocyte differentiation to macrophages is reduced and promoted when only laminin 511 is present as occurs in the *Lama4*^*-/-*^ and WT LP. Flow cytometry analyses of the main laminin binding integrins, α6 and αv, on P1, P2 and P3 populations in adult and P21 LP, revealed higher levels of integrin αv in P3 macrophages (Fig. 4F) as also observed in BMDM (Fig. S3C); however, integrin α3 was not detectable on P1, P2 or P3 populations from adult or P21 tissues.

**Fig. 4.**
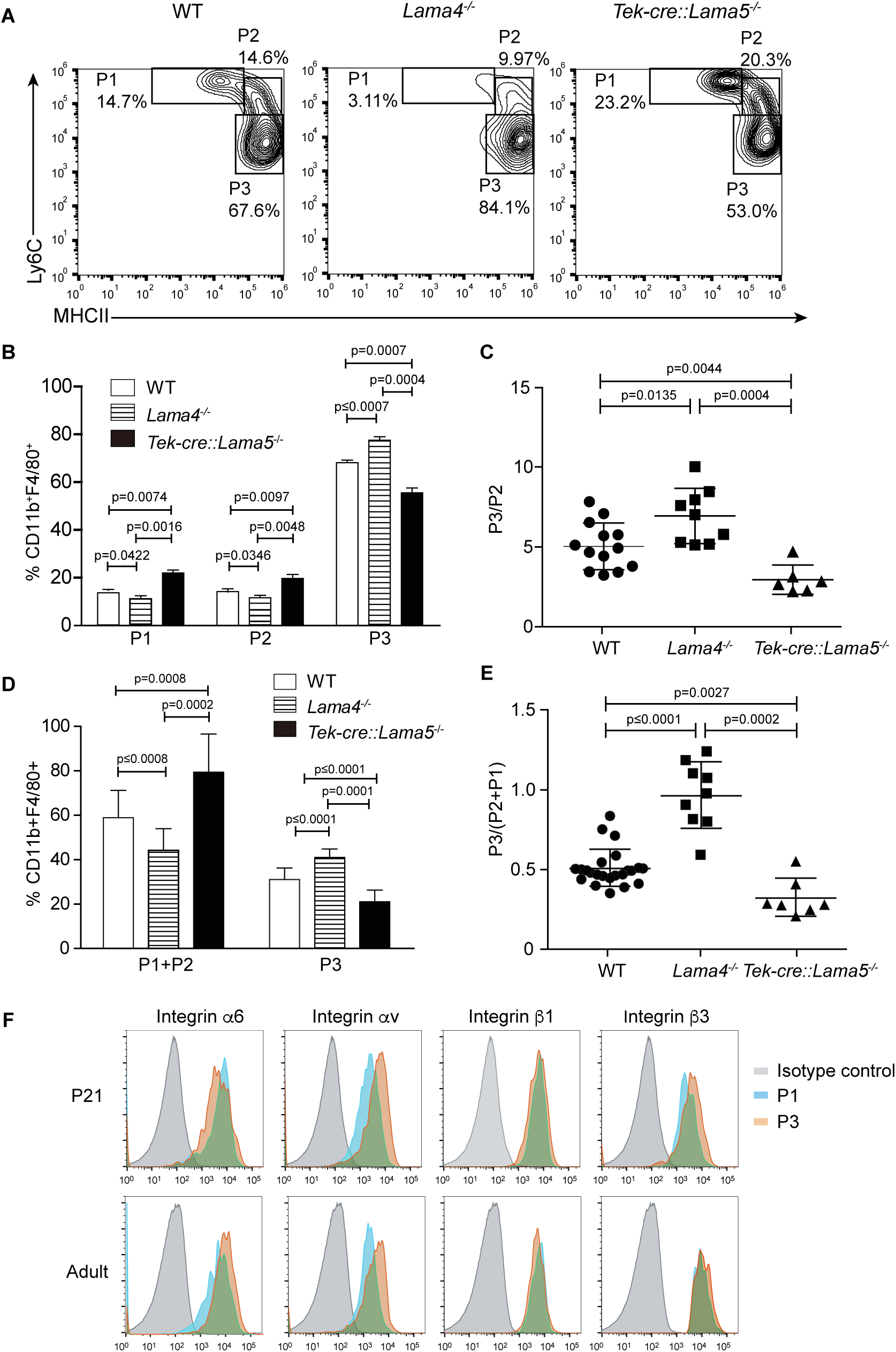
Monocyte differentiation in colons of *Lama4*^*-/-*^ and *Tek-cre::Lama5*^*-/-*^ mice. (A) Representative flow cytometry showing proportions of early extravasating P1 (Ly6C^hi^MHCII^low^) and P2 (Ly6C^mid^MHCII^mid^) populations, and mature P3 macrophages (Ly6C^low^MHCII^hi^) in isolated colonic lamina propria cells from adult WT, *Lama4*^*-/-*^ and *Tek-cre::Lama5*^*-/-*^ mice; (B) corresponding quantification of the data and (C) ratios of P3 /P2 cells. Data are means ± SD from 3 independent experiments performed with 12 WT, 6 *Lama4*^*-/-*^ and 6 *Tek-cre::Lama5*^*-/-*^ mice. (D) Corresponding analyses of P1 plus P2 populations, and mature P3 macrophages amongst colonic lamina propria cells of P21 WT, *Lama4*^*-/-*^ and *Tek-cre::Lama5*^*-/-*^ mice; and (E) ratios of P3 /P2+P1 cells. Data are means ± SEM from 3 experiments performed with at least 3 mice/ strain. Significance of the data was calculated using a Mann-Whitney test; exact *p*- values are shown in the graph. (F) Representative flow cytometry of laminin binding integrins on colon P1, P2 and P3 cells from adult and P21 WT mice. The experiment was repeat 3 times with the same pattern of results.

### Mouse splenic derived monocyte differentiation *in vitro*

As monocyte interaction with vascular endothelial was shown to promote differentiation to macrophages in an *in vitro* model (Jakubzick et al., 2013), we tested the role of laminin 511 in this setup. Endothelioma cell lines were generated from *Lama4*^*-/-*^ and WT littermate embryos (Williams et al., 1989) which expressed high (eEND4.1) or low (eENDwt) levels of laminin α5, respectively (Fig. S7A). However, despite several attempts it was not possible to establish an endothelioma cell line from *Tek-cre::Lama5*^*-/-*^ embryos as the cells did not proliferate; similar results were obtained using CRISPR/Cas 9 to eliminate *Lama5* expression in the bEND.5 cell line (manuscript in preparation). Naïve splenic monocytic cells (Fig. S7B) were incubated on confluent monolayers of eENDwt or eEND4.1 for 16 h and cells localised at the subendothelial layer were analysed by flow cytometry as described previously (Jakubzick et al., 2013), revealing a significantly higher proportion of MCHII^+^Ly6C^neg^ and F4/80^+^Ly6C^neg^ macrophages in eEND4.1 than in eENDwt cocultures (Fig. 5A, B). When the same experiments were performed with splenocytes preincubated in the presence of a function blocking antibody to integrin α6β1, the proportion of MCHII^+^Ly6C^neg^ and F4/80^+^Ly6C^neg^ macrophages in the eEND4.1 cultures were reduced to levels observed in eENDwt cultures (Fig. 5A, B). This substantiates a role for endothelial cell-derived laminin 511 in promoting monocyte differentiation to macrophages, and suggests the involvement of both the endothelium and a direct integrin α6β1-mediated interaction with laminin 511.

**Fig. 5.**
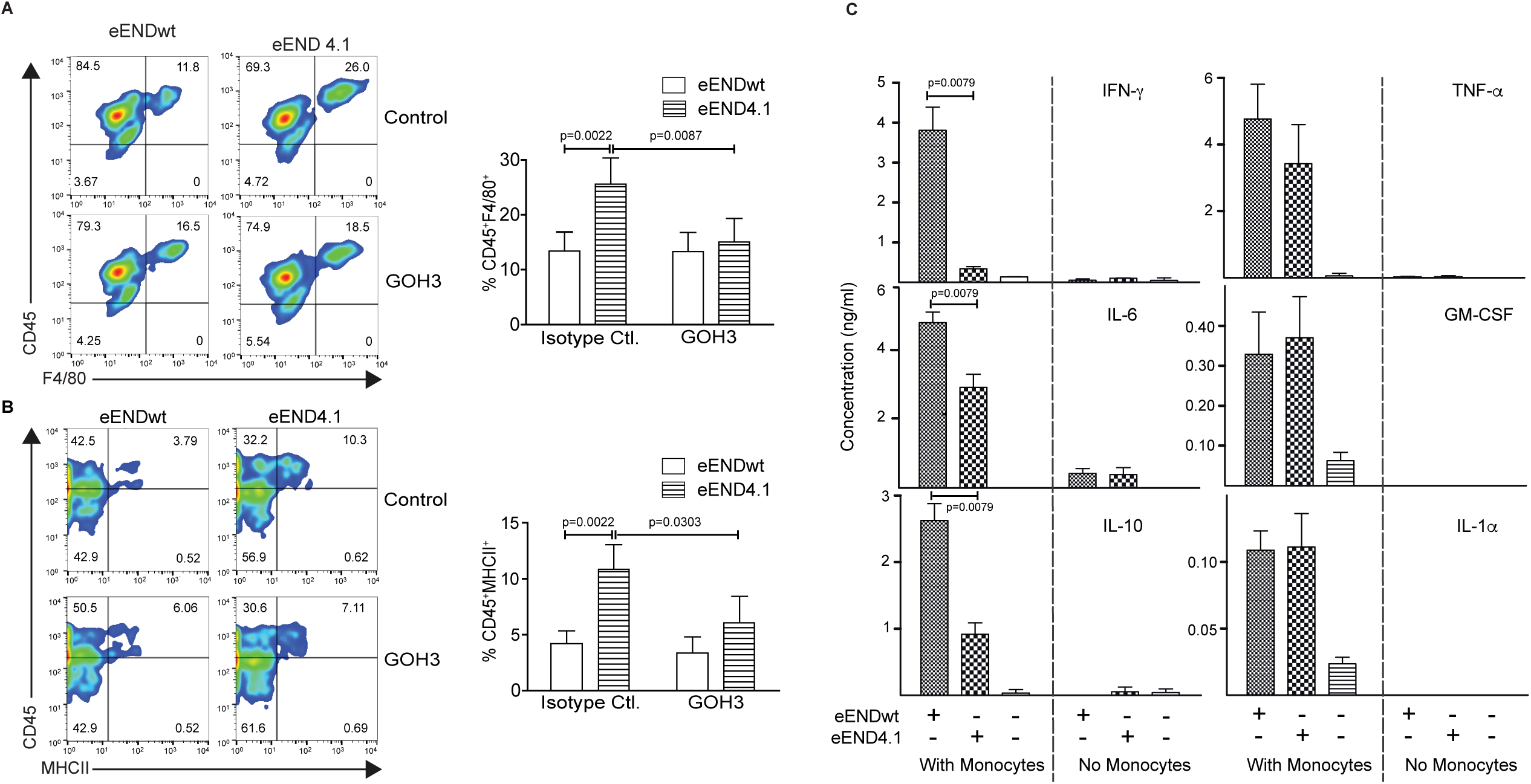
Enhanced *in vitro* differentiation of splenic monocytes to macrophages on laminin 511-overexpressing endothelial cells. Representative flow cytometry for (A) CD45^+^F4/80^+^ and (B) CD45^+^MHCII^+^ in splenic monocytes cocultured for 16h with eEND.wt or eEND4.1 cells in the presence or absence of integrin α6 blocking (GoH3) antibody; bars graphs show quantification of the data expressed as a percent of the total CD45^+^ population. Isotype control antibody had no effect. Data are means ± SD from 3 experiments with 2 replicates/experiment/treatment. Significance was calculated using a Mann-Whitney test; exact *p*-values are shown in the graph. (C) Cytokine concentrations in conditioned media of eEND.wt or eEND4.1 cells cultured in the presence or absence of splenic monocytes for 16h, and of splenic monocytes alone. Data are means ± SD of 3 experiments performed with duplicates/experiment. Absence of values indicates that the cytokines were not detectable. Significance was calculated using a Mann-Whitney test; exact *p*-values are shown in the graph.

As laminins may affect the expression of cytokines by endothelium which in turn may affect monocyte differentiation, we measured cytokine profiles in the conditioned medium of eEND4.1 and eENDwt cells cultured alone or co-cultured with the splenic monocytes, and of splenic monocytes alone under the same conditions as described above. Cytokines examined are those implicated in monocyte to macrophage differentiation, including GM-CSF, IL-6, Il-10, IFN-γ, TNF-α and IL-1α (Arango Duque and Descoteaux, 2014). Significant levels of cytokines were measured only in cocultures of eENDwt or eEND4.1 together with splenic monocytes, with lower levels of IL-6, IL-10 and IFN-γ in cocultures with eEND4.1 cells compared to eENDwt cells, but no changes in GM-CSF, TNF-α and IL-1α (Fig. 5C). Although monocytes and macrophages are the main sources of these cytokines, some such as GM-CSF, IL-6, IL-1 and IL10 can also be secreted by endothelial cells. The fact that there was no substantial secretion of these cytokines from eEND4.1 cells cultured alone suggests that laminin 511 does not affect endothelial expression of cytokines but rather that the cytokines stem from the differentiating monocytes. These data also further substantiate the hypothesis that monocyte interaction with both the endothelium and with laminin 511 are required for efficient triggering of macrophage differentiation. The pattern of cytokines observed in the eEND4.1/splenic monocyte cocultures suggests that laminin 511 promotes a non-inflammatory macrophage population (Arango Duque and Descoteaux, 2014).

## Discussion

Published data suggest that monocyte interaction with the endothelium during extravasation is a critical step in the switch to macrophage differentiation (Jakubzick et al., 2013). Our data demonstrate that it is not the endothelium alone but also laminin 511 in the underlying basement membrane that is required for this switch.

Like T cells and neutrophils, CX3CR1-GFP monocytes also preferentially extravasate at laminin 511^low^ sites, with CX3CR1-GFP^low^ inflammatory-monocytes showing preferential localization and higher motility at such sites. However, *in vitro* transmigration studies revealed an equal ability of monocytes to migrate across both endothelial laminins and non-endothelial laminins and BSA. While this may reflect shortcomings of the *in vitro* experimental systems employed, we propose that high affinity binding of the infiltrating monocytes to laminin 511 in the endothelial basement membrane traps them at this site and promotes their differentiation towards the macrophage lineage, which alters both their integrin profile and migratory mode. Both monocytes and macrophages bind efficiently to laminin 511 but only macrophages also bind to laminin 411 and can, therefore, distinguish between these two isoforms; they also migrate more efficiently on laminin 411, explaining the faster extravasation of CX3CR1-GFP cells in *Tek-cre::Lama5*^*-/-*^ hosts. This highlights the site between the endothelial monolayer and the underlying basement membrane as a critical site for the modulation of the phenotype of infiltrating monocytes and for their switch to macrophages. This is supported by the enhanced macrophage differentiation of splenic monocytes observed in cocultures with laminin 511 overexpressing endothelial cell lines and by the higher proportion of differentiating macrophages in the intestines of the *Lama4*^*-/-*^ mice, which express only laminin 511.

The adhesion of monocytes to endothelium has been reported to trigger expression of extravasation specific genes and to initiate changes towards a more differentiation phenotype (Randolph et al., 1998, Thomas-Ecker et al., 2007). More recently, specifically the site underlying endothelial cells, but not fibroblasts, was shown to be critical for this switch (Jakubzick et al., 2013), however, the molecular basis of these observations remained unclear. Our data suggest that laminin 511 together with the endothelium is critical for this step. The fact that blocking integrin α6 reduced, but did not ablate, the differentiation of monocytes incubated with endothelium overexpressing laminin 511 to macrophages suggests that two steps are involved – cross-talk with the endothelium, in an as yet undefined manner, and integrin α6β1 mediated interaction with laminin 511 in the endothelial basement membrane. The absence of such an effect in monocytes incubated with fibroblasts (Jakubzick et al., 2013) may be explained by the nature of extracellular matrix that fibroblasts secrete, which is mainly collagen types I and III and proteoglycans and, in specialized situations, also laminins 411 and 211 (Schuler and Sorokin, 1995, Frieser et al., 1997, Fleger-Weckmann et al., 2016) but not laminin 511 (Sorokin et al., 1997). The relatively long incubation time of monocytes with endothelium required for the *in vitro* differentiation to macrophages (16h), compared to the time required for monocytes to extravasate into the inflamed cremaster muscle (2-4h), suggests the absence of critical factors in the *in vitro* set up, which may include absence of a mature basement membrane that cannot be recapitulated *in vitro* using cultured cells.

There is increasing evidence that molecularly distinct steps exist subsequent to penetration of the endothelial monolayer that are critical for successful extravasation into inflamed tissues. This is suggested by intravital microscopy studies (Woodfin et al., 2011, Song et al., 2017) and by studies involving blocking of endothelial junctional molecules, such as CD99, CD99L and PECAM-1, that result in leukocyte arrested at cell-cell junctions or at the endothelial basement membrane (Liao et al., 1995, Schenkel et al., 2002, Bixel et al., 2004, Song et al., 2017, Samus et al., 2018). However, the concept that endothelial barrier function and phenotypes of extravasating immune cells may be affected by signals originating from the endothelial basement membrane, as suggested here, is only starting to emerge. How basement membrane components affect extravasation is likely to depend on the immune cell and tissue type, e.g. the effects on highly impermeable microvessels of the central nervous system are likely to differ from those on the microvessels of secondary lymphoid organs. Recent data from our lab supports a role for endothelial laminins 511 and 411 in modulating the pathogenicity of T cells infiltrating into the brain during neuroinflammation (Zhang et al., 2020). While the integrins and mechanisms involved are distinct to those described here, the data reinforce the sub-endothelial site as important for modulation of immune cell phenotypes during extravasation processes.

We show here that the switch towards a macrophage phenotype is associated with a change in integrin profile, as reported previously by others (Shaw et al., 1993, Prieto et al., 1994, Ammon et al., 2000), with the notable unique expression of integrin α3β1 by BMDM and increased surface levels of αvβ1/αvβ3 and α6β1, all of which can bind to laminin 511 and, for α6β1 and α3β1, also laminin 411 (Kikkawa et al., 2000, Fujiwara et al., 2001). Such changes are likely to affect the ability of the cells to interact with and to distinguish between the different laminin isoforms. However, differentiation towards the macrophage lineage is also associated with changes in size and cytoplasmic complexity (Jakubzick et al., 2017) that will affect migration potential and speed. The larger size of macrophages (15-20 μm) compared to monocytes (7-8 μm) may contribute to their slower migration (Hanley et al., 2010, Bzymek et al., 2016), but the associated larger cytoplasmic volume and more complex cytoskeleton may be an advantage with respect to the force required to penetrate the endothelial basement membrane *in vivo* – topics that require future investigation. Indeed, it may be serendipitous whether an extravasating monocyte encounters a laminin 511 high or low patch in the postcapillary venule basement membrane, which may determine those cells that differentiate towards the macrophage phenotype and those that do not and can re-enter the circulation (Jakubzick et al., 2017), placing endothelial laminin 511 in a pivotal position in the switch from monocytes to macrophages. While we have here specifically examined monocytes/ macrophages, the processes and concepts described may be relevant to other differentiation processes involving cells located at such vascular niches, including stem cells.

## Acknowledgements

We thank Sigmund Budny for purification of laminins, Barbara Liel for generation of eENDwt and eEND4.1 cells, and Hanna Gerwien for all statistical analyses. This work was supported by funding to LS from the German Research Foundation (CRC1009 A02, EXE1003).

## Contribution to the field

The manuscript deals with the differentiation switch of monocytes to macrophages that occurs during extravasation, which is known to critically require monocyte interaction with the endothelium; however, the molecular mechanism/s involved remain unclear. We show that it is not the endothelium alone but also the underlying basement membrane that is critical for this switch and identify laminin 511, an integral endothelial basement membrane component, as a central player in this process. Most importantly, our work highlights the subendothelial site as a site of immune cell modulation, which in turn influences the ability of immune cells to penetrate the basement membrane barrier and infiltrate into the inflamed tissue. This is important fundamental information relevant to most, if not all, inflammatory processes and also to differentiation processes involving other cell types located at such vascular niches.

## Author contributions

Lixia Li carried out the intravital microscopy (IVM) experiments and most adhesion and migration assays; Jian Song analysed all IVM experiments and carried out all *in vitro* macrophage differentiation experiments; Sophie Loismann performed adhesion assays with WT and CD18^-/-^ Hoxb8 monocytes; Omar Chuquisana performed flow cytometry of Hoxb8 monocytes and BMDM; Thomas Vogl generated WT and CD18^-/-^ Hoxb8 monocyte lines and contributed to writing of the manuscript; Johannes Roth contributed to experimental design and writing of the manuscript; Rupert Hallmann generated and characterised endothelioma cell lines and, together with Melanie-Jane Hannocks, contributed to experimental design, analyses of data and writing of the manuscript; Lydia Sorokin designed the project, supervised all work and wrote the original manuscript.

