## Supplementary material for "Endothelial basement membrane laminins as an environmental cue in monocyte differentiation to macrophages": Supplemnetary Figures

**Supplemental Information**

**Fig. S1 Phase contrast and fluorescence images taken from movies of CX3CR1-GFP$^+$ cells extravasating across postcapillary venules of the cremaster muscle of WT, *Lama4$^{-/-}$* and *Tek-cre::Lama5$^{-/-}$* mice.** Boxed areas are 75 μm from either side of the postcapillary venule for a length of 100 μm, that were used to quantify extravasated CX3CR1-GFP$^+$ cells; scale bars = 50μm. The first (WT) column is also shown in Fig. 1A.

**Fig. S2 In situ cytometry of CX3CR1-GFP$^+$ mean fluorescence intensity (MFI) at laminin α5 low (left) and high (right) sites during CCL-2 induced extravasation across postcapillary venules of WT mice and correlation between speed of migration of individual cells and GFP+ MFI.** (A) To track CX3CR1-GFP$^{low}$ inflammatory monocytes GFP mean fluorescence intensity (MFI) was measured *in situ* in areas of low and high laminin 511 expression, revealing a higher proportion of CX3CR1-GFP$^{low}$ cells at laminin 511$^{low}$ (left) compared to laminin 511$^{high}$ sites (right). Quantification of this data is shown in Fig. 2B. (B) Representative plot of the correlation between GFP MFI (Y axis, left, blue line) and the migration speed (Y axis, right, green dots) of 80 individual cells (X-axis) from 39 CX3CR1$^{hi}$ and 41 CX3CR1$^{low}$ sites analysed in 3 WT hosts. Quantification of this data is shown in Fig. 2C.

**Fig. S3 Representative flow cytometry for integrin subunits.** (A) Primary human monocytes, (B) mouse monocyte-like Hoxb8 cells derived from CD18$^{-/-}$ mice and their WT littermates, and (C) WT bone marrow derived macrophages (C).

**Fig. S4** Transmigration of human monocytes (A) and WT monocyte-like Hoxb8 cells (B) across HUVEC or mouse bEND.5 cells plated on laminins 111, 411, or 511 pre-coated transwell inserts. Transmigrated cells were expressed as percentage of total cells added. Data are means ± SD from 3 (B) or 4 (A) independent experiments with three replicates/experiment/treatment.

**Fig. S5 Dot blots for the expression of laminin α4 and laminin α5 polypeptides in mouse and human monocytic cells and sera; related to Figure 4.** Cell lysates from untreated and LPS treated mouse Hoxb8 precursors and monocytes, bone marrow derived macrophages (BMDM), and sera from WT and *Lama4$^{-/-}$* mice were analysed by dot-blot using rabbit anti-mouse laminin α4 antibody (377) (Ringelmann et al., 1999), pre-immune serum and secondary antibody only. (B) Mouse Hoxb8 monocytes and BMDM were analysed using purified rabbit anti-mouse laminin α5 (405) (Ringelmann et al., 1999). (C) Human monocyte lysates and sera were blotted and stained with mouse anti-human laminin α4 (3D12) (Korpos et al., 2013) or only 2$^{nd}$ antibody. (D) Human monocyte lysates were analysed using mouse anti-human laminin α5 (6A11) (Korpos et al., 2013). Dots of 5μg total protein or serum were analysed; dashed lines mark area of drops.

**Fig. S6 Gating strategy employed for identification of P1, P2 and P2 differentiating macrophage populations in the colon lamina propria.** Modified from Bain et al., (2014)(Bain et al., 2014). Cell aggregates and dead cells were excluded by FSC and viability dye staining, respectively. Total leukocytes were selected by CD45 expression; Ly6G$^+$

neutrophils and Siglec-F$^{+}$ eosinophils were then gated out from CD45$^{+}$ population. To obtain F4/80$^{+}$CD11b$^{+}$ cells, the CD11c$^{high}$F4/80$^{low}$ mucosal dendritic cells (DC), were gated out. Based on Ly6C and MHCII levels, the F4/80$^{+}$CD11b$^{+}$ cells were further divided to P1 (Ly6C$^{high}$MHCII$^{low}$), P2 (Ly6C$^{mid}$MHCII$^{mid}$) and P3 (Ly6C$^{low}$MHCII$^{high}$) populations. (B) representative flow cytometry of P21 colons for P1 plus P2 and P3 populations. Quanitification of this data is shown in Fig. 4D, E.

**Fig. S7 Characterization of eEND4.1 and eENDwt endothelioma cell lines.** Endotheliomas were derived from *Lama4*$^{-/-}$ embryos (eEND4.1) and their WT littermates (eENDwt) and flow cytometry of splenic monocytes employed in experiments shown in Fig. 7 of the main text. (A) Full Western blot with secondary antibody control and tubulin control shown. Representative flow cytometry of F4/80 (B) and MHCII (C) on sorted splenic monocytes before (left) and after (right) 16 h incubation in the absence of endothelial cells.

**Movie 1** Phase contrast intravital imaging of CCL-2 induced monocyte extravasation in a cremaster muscle model performed in a WT mouse, showing rolling and adherent cells within the vessel lumen, as well as cell penetrating into the surrounding tissues.

**Movie 2** Fluorescent imaging of CX3CR1-GFP+ cells and laminin $\alpha5$ in the endothelial basement membrane (left) and of laminin $\alpha5$ alone (right) during CCL2-induced monocyte extravasation. GFP+ cells are pseudo-colored, with higher levels of GFP occurring in the warmer colors and lower levels in the cooler colors; arrows mark sites of lower laminin $\alpha5$ expression where extravasation preferentially occurs.
